# Evolution of Conditional Cooperativity Between HOXA11 and FOXO1 Through Allosteric Regulation

**DOI:** 10.1101/014381

**Authors:** Mauris C. Nnamani, Soumya Ganguly, Vincent J. Lynch, Laura S. Mizoue, Yingchun Tong, Heather L. Darling, Monika Fuxreiter, Jens Meiler, Günter P. Wagner

**Affiliations:** Yale Systems Biology Institute and Department of Ecology and Evolutionary Biology, Yale University, New Haven, CT, 06511 USA; Departments of Chemistry, Pharmacology and Biomedical Informatics; Center for Structural Biology and Institute of Chemical Biology, Vanderbilt University, TN 37232, USA; Department of Human Genetics, The University of Chicago, 920 E. 58th Street, CLSC 319C, Chicago, IL 60637 USA; Howard Hughes Medical Institute, Department of Chemistry and Biochemistry, University of Colorado, Boulder, CO 80303; MTA-DE Momentum Laboratory of Protein Dynamics, University of Debrecen, H-4032 Debrecen, Nagyerdei krt 98, Hungary

## Abstract

Transcription factors (TFs) play multiple roles in different cells and stages of development. Given this multitude of functional roles it has been assumed that TFs are evolutionarily highly constrained. Here we investigate the molecular mechanisms for the origin of a derived functional interaction between two TFs that play a key role in mammalian pregnancy, HOXA11 and FOXO1. We have previously shown that the regulatory role of HOXA11 in mammalian endometrial stromal cells requires an interaction with FOXO1, and that the physical interaction between these proteins evolved long before their functional cooperativity. Through a combination of functional, biochemical, and structural approaches, we demonstrate that the derived functional cooperativity between HOXA11 and FOXO1 is due to derived allosteric regulation of HOXA11 by FOXO1. This study shows that TF function can evolve through changes affecting the functional output of a pre-existing protein complex.

## Introduction

Most TFs play roles in different tissues, cells, and developmental stages (Hu and Gallo, 2010). For instance HOXA11 is involved in the development of limb, kidney, male and female reproductive tracts, cloaca and hindgut, and in the function of T- and B-cells (Davis et al., 1995; Hsieh-Li et al., 1995; Schwab et al., 2006; Speleman et al., 2005; Wellik and Capecchi, 2003; Yokouchi et al., 1995). Given its many pleiotropic roles, one might expect random mutations in HOXA11 to have a high likelihood to disrupt at least one of its many functions. However, there is evidence that the HOXA11 protein underwent functionally advantageous changes in the stem lineage of placental mammals (Lynch et al., 2008). Evolutionary changes to TF function have been documented before, e.g. *Tinman/Nkx2.5* (Ranganayakulu et al., 1998; Schwartz and Olson, 1999), *Ubx* (Galant and Carroll, 2002; Grenier and Carroll, 2000), flower development genes (Bartlett and Whipple, 2013; Lamb and Irish, 2003), *HOM/Ftz* (Lohr et al., 2001); and many more (Wagner and Lynch, 2008). These examples highlight a discrepancy between the model of conserved TF genes function and the empirical facts documenting evolutionary changes in TF function. A well-documented mode of TF protein evolution is the acquisition of Short Linear Motifs (SLiM) (Diella et al., 2008; Neduva and Russell, 2006). In this model of TF evolution, new functionalities arise via derived protein-protein interactions that are usually embedded in a protein segment which lack a well-defined three-dimensional structure (Fuxreiter et al., 2007). Here we provide evidence for another mechanism of TF evolution: the evolution of intra-molecular regulation within the HOXA11 protein resulting in a novel functional output in the presence of a pre-existing TF partner, FOXO1.

Endometrial stromal cells are part of the inner lining of the uterus and play an important role in mammalian pregnancy. To accommodate the implantation of the conceptus, the stromal cells differentiate into the so-called “decidual cells.” Decidual cells express a number of genes that are important for the maintenance of pregnancy, most notably decidual prolactin (dPRL) and IGFBP1(Kutsukake et al., 2007; Tseng and Mazella, 2002). HOXA11 plays a critical role in the expression of these decidual genes (Lynch et al., 2009). In eutherian mammals, HOXA11 cooperates with FOXO1 to induce expression of dPRL, while HOXA11 alone is a repressor (Lynch et al., 2009; Roth et al., 2005).

In this paper we demonstrate that the activator function of HOXA11 was already present in the common ancestor of mammalian HOXA11 proteins. In the absence of FOXO1, however, the activation function of Eutherian HOXA11 proteins is repressed through an intra-molecular interaction. We show that evolutionary changes to the HOXA11 protein led to FOXO1 dependent unmasking of the intrinsic transcriptional activation function of HOXA11.

## Results

Our model for decidual gene regulation is the decidual prolactin promoter active in human decidual cells (Berwaer et al., 1994; Gellersen et al., 1994; Gerlo et al., 2006). Three findings from our previous work guide our experimental approach: 1) the amino acid substitutions in HOXA11 protein are all located amino (N)-terminal to the homeodomain (Lynch et al., 2008); 2) The physical interaction between HOXA11 and FOXO1 arose before the functional cooperativity (Brayer et al., 2011); 3) The derived cooperativity is due to evolutionary changes in HOXA11 (Brayer et al., 2011; Lynch et al., 2008). There is no structural information about the N-terminal region of HOXA11 that could guide our experimental analysis. We first conducted a computational structural biology analysis to determine sequence segments, which could embed functional motifs.

### HOXA11 N-terminus is disordered with a tendency to form α-helices

The multiple sequence alignment of HOXA11 sequences from different species indicates highly conserved amino acid residues in regions 1-67 and 125-151. Large variation is seen in the range from amino acid 68-124 and 152-173. Beyond amino acid 173 large gaps, low sequence conservation, and multiple repeats are found up to amino acid 280 indicating potentially disordered region (Figure 1A&B). Secondary structure predictions using PSIPRED (Buchan et al., 2010; Jones, 1999) and JUFO (Leman et al., 2013) predict several short peptides with a tendency to form α-helices (AAs 85-93, 104-107 and 140-148) or β-strands (around AAs 14, 20, 44, and 60, Figure 1C). We hypothesized that these segments correspond to regions of intrinsic disorder that might form structure when interacting with partner proteins (Buchan et al., 2010; Dyson and Wright, 2005; Fuxreiter et al., 2004; Jensen et al., 2009; Jones, 1999; Leman et al., 2013; Tompa, 2005; Vucetic et al., 2005). To test if formation of secondary or tertiary structure is plausible we utilized the *de novo* protein structure prediction algorithm Rosetta (Das et al., 2009; Fleishman et al., 2011; Simons et al., 1997). We folded residues 1-150, 58-155, and 85-157 in three independent experiments. In all cases Rosetta introduced α-helices and β-strands in the regions that were predicted to have some tendency to form secondary structure. Furthermore, Rosetta sometimes folded regions 58-154, and 85-157 into helical bundles consisting of three to four α-helices (Figure 2A & B). We concluded that region 58-157 has a tendency to form α-helical secondary structure. We named the region 64-152 IDR and region 80-152 ΔNP-IDR (Suppl. Table 1).

**Figure 1.**
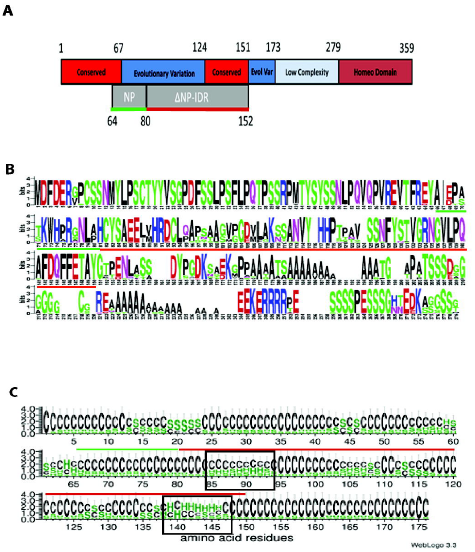
(**A**) Schematic representation of HOXA11 showing the overall domain organization and the sequence variability along different regions. The regions with conserved sequences (red), larger evolutionary variation (blue), low sequence complexity (light blue) and highly conserved homeodomain (brown) were identified from multiple sequence alignment of HOXA11 protein sequences from several different species. The boxes in gray indicate the two regulatory domains between regions 64-152 (IDR) identified during this study, which include the negative regulatory domain (NP, green underlined) and the disordered regulatory region (ΔNP-IDR, red underlined). (**B**) Sequence logo of HOXA11 amino acid sequence generated using weblogo after multiple sequence alignment. The height of each amino acid is proportional to the degree conservation of residue in a given position of the sequence. Regions for the NP and ΔNP-IDR are indicated with red and green underlines respectively. (**C**) Weblogo representation of *HOXA11* IDR secondary structure prediction. The secondary structure for *HOXA11* residues 1-175 was predicted using jufo. The vertical axis denotes the probability of a coil (C), Helix (H) or a strand (S) to occur for a given residue represented in the horizontal axes. The most prominent helix structures are shown in the black box. Regions for the NP and ΔNP-IDR are indicated with red and green overhead lines respectively.

**Figure 2.**
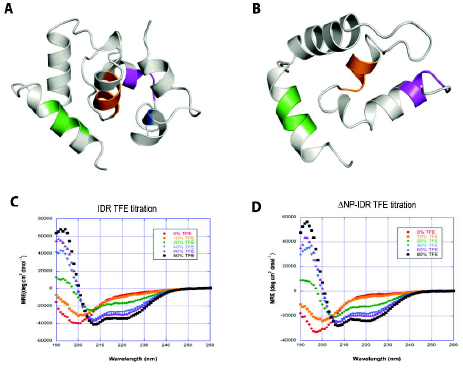
Denovo models of HOXA11 structure predicted by ROSETTA for residues (**A**) 58-154 IDR and (**B**) residues 85-159 (ΔNP-IdR). The colors indicate three putative KIX binding domains at position 85-88 (magenta), 103-107 (orange, also called PIM for Protein Interaction Motif in the paper), 142-146 (Daniels et al.). The single tryptophan residue in IDR at position 73 is colored blue. Circular Dichroism spectra showing the effect of increasing TFE concentration on the helical propensity of (**C**) IDR and (**D**) ΔNP-IDR. Increasing concentration of TFE led to 35% helicity in both HOXA11 constructs.

To test secondary structure content experimentally, circular dichroism spectra (Macdonald et al., 1964) were collected for the hOxA11 constructs IDR (64-152) and a ΔNP-IDR (80-152). The spectra were similar for both constructs and exhibited characteristics of intrinsically disordered proteins dominated by a strong negative band at 200nm with shallow minima at 222nm indicative of a small amount of α-helical structure. Addition of 2,2,2-trifluoroethanol (TFE) caused rapid increase in α-helicity for both constructs from around 8% to 35% (Figure 2C & D). We conclude that little secondary structure is present in aqueous solution but α-helical character can be readily induced.

### The intrinsically disordered region is critical for regulatory function

To determine the functional role of the IDR, we designed eight N-terminal truncation mutants of HOXA11 (Figure 3A) (Roth et al., 2005) and tested their ability to trans-activate luciferase expression from the dPRL promoter when co-expressed with FOXO1 (Figure 3B). We found that co-transfection of wild type HOXA11 and FOXO1 resulted in an up-regulation of luciferase expression, confirming our previously reported results (Lynch et al., 2009; Lynch et al., 2008) (Figure 3C). Truncation of the HOXA11 protein up to amino acid 130 (ΔN130) had a significant negative effect on the cooperative up-regulation of luciferase expression from the dPRL promoter, but still retained significantly higher expression compared to background (t-test p < 0.001), whereas a truncation to amino acid 150 (ΔN150) resulted in a complete loss of reporter gene activation. Further truncations of the protein restored reporter gene expression. Unexpectedly an internal deletion of the predicted IDR region (Δ66-151) did not negatively impact trans-activation (Figure 3C). Collectively these results suggest that the N-terminal region of HOXA11 is a multifunctional disordered segment, which contains critical intramolecular regulatory sites/motifs that modulate the trans-activation functions of HOXA11.

**Figure 3.**
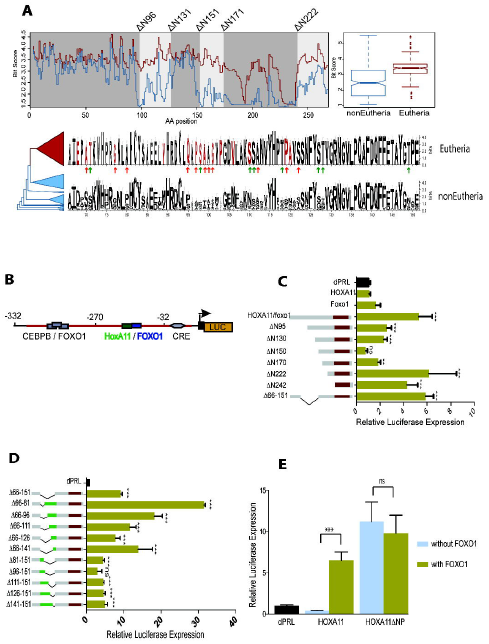
Characterization of the Intrinsically Disordered Region (IDR). **A** Conservation plot of Eutherian (n=82, red line) and non-Eutherian (n=55, blue line) HOXA-11 proteins. **A**, (bottom), Sequence conservation of N-terminal amino acids of HOXA11. Derived Eutherian amino acid changes are shown in red. **B**, Diagram of the luciferase reporter vector and experimentally characterized transcription-factor binding sites. **C**, Gene reporter assays of N-terminal deletion constructs of Eutherian HOXA11 (mouse) co-transfected with FOXO1 in human ESCs (HESC) cell lines. Results suggest HOXA11 N-terminal is a multifunctional domain. **D**, Detailed deletion constructs of the IDR revealed the NP (residues 6681) as a critical region of the IDR required for regulatory cooperativity. **E**, ΔNP HOXA11 is FOXO1 independent in transactivation activity. Luciferase values are shown as fold changes (mean ± s.e.m., n = 6) relative to the reporter control (dPRL). *p < 0.05, **p < 0.01, ***p < 0.001, ns p > 0.05.

### Minimal region of the IDR required for regulatory cooperativity

We performed a detailed deletion scan of the IDR to further characterize its role in regulating the trans-activation functions of HOXA11. We generated five internal deletions from the N-terminus of the IDR (ΔN66-81, ΔN66-96, ΔN66-111, ΔN66-126, ΔN66-141) and five internal deletions from the C-terminus of the IDR (ΔC141-151, ΔC126-151, ΔC111-151, ΔC96-151, ΔC81-151) and tested their ability to trans-activate reporter genes. The most notable deletion construct is HOXA11 Δ66-81, which showed the strongest enhancement (>10x) of activity relative to wild type HOXA11 (Figure 3D).

The trans-activation abilities of the N-terminal deletions suggest the presence of a regulatory region within amino acids 66-81 with a strong repressive effect on the activation function of HOXA11 (Figure 3D). We call the region 66-81 “Negative Regulatory Peptide” (NP). We hypothesize that the NP masks an activation domain. Moving forward, we refer to IDR construct 80-152 as ΔNP-IDR (deleting residues 66-81).

### The ΔNP HOXA11 mutation is a FOXO1 independent activator

We hypothesized that the NP is an intra-molecular repressor of HOXA11 activation function and that the interaction between HOXA11 and FOXO1 is relieving the repressive effect of the NP. This model predicts that the deletion of the NP fragment, ΔNP, should lead to FOXO1 independent activation. We tested this prediction and found that the HOXA11ΔNP construct trans-activated luciferase expression independently of FOXO1 (Figure 3E). These results are consistent with an intra-molecular regulatory role of the NP and suggest that the role of FOXO1 interaction is to relieve the repressive effect of the NP.

### The NP regulatory peptide is ancestral to Therians

We have previously resurrected the ancestral therian HOXA11 protein (AncThHOXA11), the protein present in the last common ancestor of placental mammals (Eutheria) and marsupials. We found that the resurrected protein was expressed, localized to the nucleus, and appropriately regulated target genes (Brayer et al., 2011; Lynch et al., 2008). The ancestral therian HOXA11 was shown to have the same repressive effect as the eutherian HOXA11 but is unable to cooperatively up-regulate dPRL expression in the presence of FOXO1.

We produced and assayed an NP deletion construct of the AncThHOXA11 (i.e. Δ66-81, AncThA11ΔNP). The AncThA11ΔNP construct showed an increase in activation similar to the eutherian HOXA11ΔNP and was also found to be FOXO1 independent. (Figure 4A). These results indicate that the regulatory function of the NP is an ancient feature of HOXA11 that predates the divergence of marsupials and eutherians, and that the eutherian protein evolved the ability to relieve the repression by the NP by interaction with FOXO1.

**Figure 4.**
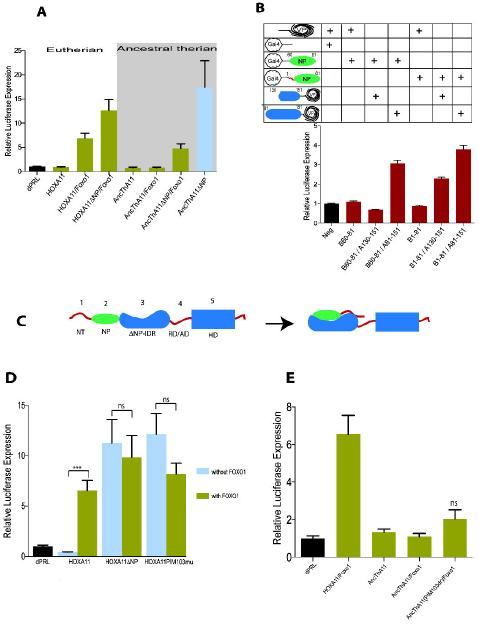
Functional equivalence of the NP between Eutherian and Ancestral Therian HOXA11. **A**, Reconstructed ancestral therian (AncThA11) ΔNP mutant construct (gray shaded area) showed similar repressive function to eutherian HOXA11 (non-shaded area). The ΔNP Ancth-A11 also showed FOXO1 independence in transactivation function (blue column). **B**, Mammalian Two Hybrid (M2H) assay identifies interaction between the NP and IDR (prey). Luciferase values are shown as fold changes (mean ± s.e.m., n = 6) relative to background measured using CheckMate Negative Control Vectors: pBIND and pACT. **C**, Schematic illustration for the intramolecular interaction between the IDR and N-terminal region. **D**, Both ΔNP and backward mutation of derived PIM103 site in the eutherian HOXA11, are FOXO1 independent transactivators. **E**, Forward mutation of the derived PIM 103 in ancestral therian HOXA11 was not sufficient in recovering cooperative transactivation activity.

### The NP has intra-molecular interactions

The functional assays we described above suggest that the NP masks the ability of HOXA11 to trans-activate, which is relieved upon interaction with FOXO1. To more directly test this hypothesis we used a mammalian-two-hybrid (M2H) system to detect physical interactions between the NP (60-81), an extended-NP (1-82), ΔNP-IDR (81-151), short-IDR (130-151). We found that co-transfection of the NP construct with the ΔNP-IDR construct, but not with the short-IDR construct, led to a significant increase in luciferase expression. In contrast cells co-transfected with the extended-NP and either the short-IDR or the ΔNP-IDR, lead to an increase in luciferase expression (Figure 4B). These data suggest that NP physically interacts with residues 82-130 of the IDR, whereas residues 1-59 region interacts with residues 130-151 of the IDR (Figure 4C).

### A derived putative interaction motif interacts with NP in preventing activation

Our M2H results suggest that the NP interacts with residues between amino acids 81 and 130 (Figure 4B and C). To identify a putative interaction site we looked for evolutionarily derived protein-protein interaction motifs, i.e. regions that 1) have a derived amino acid sequence in the eutherian HOXA11, i.e different in the ancestral HOXA11 protein, 2) have a tendency to form α-helices that can serve as scaffold for the interaction, and 3) contain changes in the pattern of Prolines or Glycines between ancestral and eutherian HOXA11 as Prolines or Glycines are known to change α-helical propensity (Cordes et al., 2002; Jacob et al., 1999). Interestingly, at AA 103-107 we found a sequence, PGDVL, derived in the eutherian HOXA11 of interest as P103 is not present in the ancestral HOXA11 protein and residues 104-107 have a tendency to form α-helcies accorindg to our secondary structure prediction (Figure 2A &B). We label this region as Putative Interaction Motive (PIM).

To test whether the derived PIM is responsible for the intra-molecular interaction with the NP, we introduced two back mutations in the eutherian HOXA11 protein to their state in the ancestral therian protein (PIM103mu; i.e. AA’s 103-107 PGDVL to GDML, P103 deletion and V106M substitution), and performed reporter gene assays as previously described. As expected, the PIM103mu mutant trans-activated luciferase expression from the dPRL reporter vector independent of FOXO1, similar to the ΔNP construct (Figure 4D). We then introduced the derived P103 and M106V substitutions into the AncThHOXA11 construct (PIM103dr) to determine if these changes were sufficient to induce cooperativity with FOXO1 (Figure 4E). Although we observed a slight increase in transactivation activity, the change was not significant and could not explain the cooperative up-regulation seen with the eutherian HOXA11 protein. These results suggest that the derived PIM103 site is necessary but not sufficient for the cooperative interaction with FOXO1.

### CBP contributes to cooperative transcriptional regulation

The data presented above suggest a phenomenological model of how cooperative target gene activation is achieved through the interaction between eutherian HOXA11 and FOXO1 proteins. To identify the mechanistic underpinnings we first identified potential co-factors that mediate target gene activation. The histone acetyltransferase CREB-binding protein, CBP, is an activating cofactor for many HOX proteins (Bei et al., 2007; Chariot et al., 1999; Choe et al., 2009). The majority of TF-CBP interactions are mediated through the KIX binding domains (KBD) that interacts with peptides that have the φ-x-x-φ-φ motif (φ is a large hydrophobic residue and “x” is any residue) (De Guzman et al., 2006; Lee et al., 2009; Plevin et al., 2005; Radhakrishnan et al., 1997; Wang et al., 2009).

Analysis of HOXA11 revealed seven sequences with similarity to the KBD (Suppl. Table 2). Of these three are located in the IDR with only one following the sequence pattern perfectly. The “perfect KBD” (residues142-146), is predicted to have a strong helical propensity by PSIPRED, JUFO and by ANCHOR calculations (Figure S1B) (Meszaros et al., 2009).

To determine the functional role of CBP in the cooperative transactivation of dPRL by HOXA11 and FOXO1A, we co-transfected CBP, HOXA11 and FOXO1A and tested their ability to trans-activate gene expression. Up-regulation of the reporter gene was more pronounced with the addition of the CBP expression vector compared to transfections with HoxA11 and FOXO1 alone (Figure 5A). This result suggests that CBP contributes to target gene activation by the HOXA11-FOXO1A complex.

**Figure 5.**
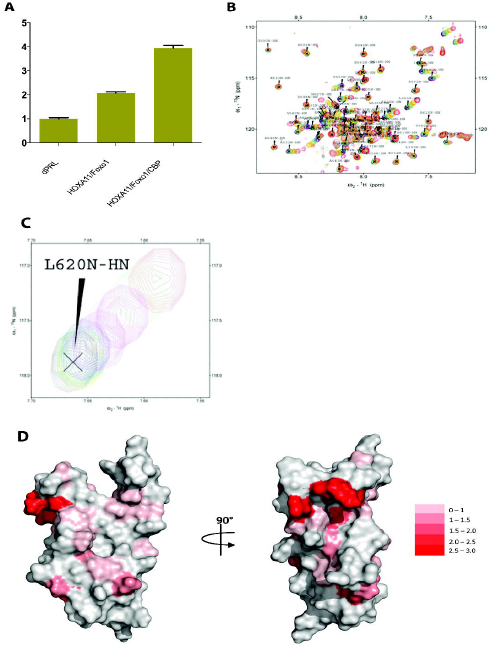
The role of CBP in HOXA11/FOXO1 dependent gene activation. **A**, Cooperative transactivation activity between HOXA11 and FOXO1A is increased by CBP. Gene reporter assay was performed for Hela cells co-transfected with HOXA11, FOXO1A, and CbP. Results suggest that the addition of CBP increases cooperative transactivation off the dPRL that has been previously described for HOXA11 and FOXO1A. Luciferase values are shown as fold changes (mean ± s.e.m., n = 6) relative to the reporter control (dPRL). *p < 0.05, **p < 0.01, ***p < 0.001, ns p > 0.05. **B,** Overlaid ^1^H-^15^N HSQC spectra of ^15^N labeled KIX (40μM) titrated with up to 20 molar equivalent of unlabeled ΔNP-IDR. **C**, Expanded region of the NMR spectra showing the chemical shift changes of residue L620 of KIX with increasing ΔNP-IDR concentration. **D**, Normalized backbone chemical shift changes of KIX up on titration of 20 molar equivalent of ΔNP-IDR mapped on the solution structure of KIX-MLL-cMyb ternary complex (PDB: 2AGH). Residues marked with different shades of red indicates standard deviation from the average chemical shift change of 0.04 ppm. The sites corresponding to cMyb (left) and MLL (right) are presented through a 90° rotation along the vertical axes.

### KBD142-146 of ΔNP-IDR binds with KIX domain of CBP

The above results suggest that CBP contributes to target gene activation. We hypothesized that KBD142-146 mediates binding of CBP. In order to test this model we investigated the interactions between 80-152 (ΔNP-IDR) and the KIX domain (residues 586-672 of mouse CBP). The ΔNP-IDR HOXA11 construct was titrated into ^15^N labeled KIX domain, and corresponding ^1^H – ^15^N heteronuclear single quantum coherence (HSQC) spectra were recorded (Figure 5B & C). Normalized chemical shift changes for each backbone amide group upon the addition of ΔNP-IDR were calculated. Chemical shift differences of 0.04ppm or higher were considered to be significant. Published NMR assignments for KIX (Radhakrishnan et al., 1997) were used to map the binding interface of HOXA11ΔNP-IDR. Fast chemical exchange was observed between the bound and unbound states of KIX. Significant chemical shift changes were observed for residues F612, T614, L620, K621, M625, E626 and N627 corresponding to the MLL binding site of KIX (Goto et al., 2002), and residues L607, Y650, H651, I660 and E665 correspond to the cMyb binding site of KIX (Zor et al., 2002) (Figure 5D & Figure S2A). The overall binding affinity of KIX was quite low (∼0.5±0.02mM) indicating a weak interaction common for disordered proteins interacting with transcription activators (Wang et al., 2012). Analysis of the two KIX binding pockets revealed that in fact the binding affinity is two times higher at the MLL site (K_d_ ∼0.33±0.02mM) than at the cMyb site (Figure S2B).

Reverse titration of KIX with ^15^N labeled ΔNP-IDR produced further insight into the structural characteristics of this region when bound to KIX (Figure S3). ^1^H-^15^N HSQC spectrum of ΔNP-IDR shows low signal dispersion in the ^1^H dimension suggesting an intrinsically disordered state even when bound to KIX (Figure S3A&B). Significant changes in chemical shift differences were observed for at least eleven ΔNP-IDR residues. Because of the low signal dispersion and large number of proline residues embedded within the sequence, we were only able to partially assign ΔNP-IDR residues. The peaks exhibiting significant chemical shift changes were assigned to those corresponding to residues 142-146, i.e. the putative “perfect” KBD, and a few residues flanking the motif.

To further test whether the KBD 142-146 is necessary for the interaction with the KIX domain we mutated the hydrophobic residues at 142-146 (ΔNP-IDR 142-146) (FDQFF to ADQAA). These mutations resulted in complete loss of binding with no significant chemical shift changes observed indicating the importance of these residues for the binding of the KIX domain and the recruitment of CBP (Figure S3C). We conclude that the HOXA11-FOXO1A complex drives target gene expression, at least in part, through the recruitment of CBP to KBD142-146.

### HOXA11 evolved DNA-pk kinase phosphorylation sites necessary for transactivation

Mass-spectroscopy results suggest that HOXA11 is differentially phosphorylated upon hormone stimulation; therefore, phosphorylation might play a role in HOXA11 activation function (Figure S4 and Suppl. Table 3). We investigated which kinases are responsible for regulating HOXA11 functional activity. Two of the identified sites S98 and T119, were derived phosphorylation sites in eutherian mammals. Computational analysis identified 6 kinases predicted to phosphorylate S98 or T119 (Figure 6A and Suppl. Table 4). We then used transcriptome data to identify which of these kinases were expressed in hESCs. We found that the kinases ERK1/2, GSK-3, DNA-pk, and CDK 2/5 were predicted to phosphorylate S98 or T119 and were highly expressed in hESC. To test if phosphorylation by these kinases played a role in potentiating transactivation by HOXA11 we performed luciferase reporter assays with eutherian HOXA11 and FOXO1, but blocked kinase activity with either ERK1 Inhibitor II, GS-3 Inhibitor XIII, DNA-pk inhibitor III, or CDK 2/5 kinase inhibitors. We found that blocking either GSK-3 or DNA-pk inhibited transactivation from the dPRL reporter vector (Figure 6B).

**Figure 6.**
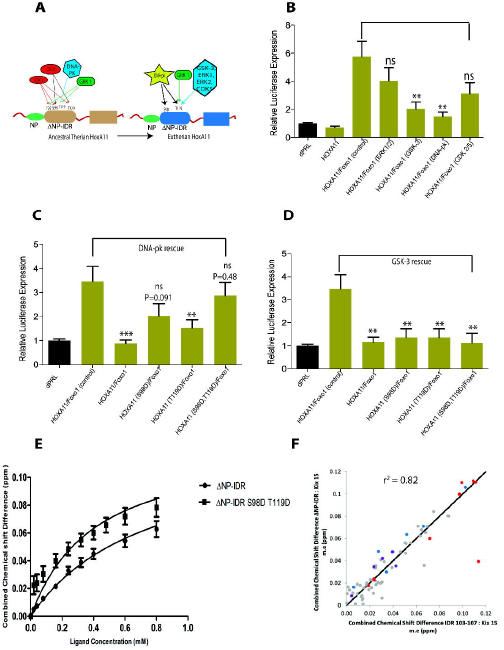
Gain in function of HOXA11 is mediated by DNA-PK kinase activity. **A**, Schematic illustration of the gain and lose of Kinase motifs between ancestral therian (left image) and eutherian HOXA11 (right image). **B**, Effects on the cooperative transactivation activity of HOXA11 and FOXO1 in HESCs treated with ERK1/2, GSK-3β, DNA-pk, and CDK 2/5 kinase inhibitors. Rescue of inhibition on transactivation activity by kinase inhibitors in (**C**) DNA-pk and (**D**) GSK-3ß by phospho-simulation at amino acids S98D and T119D. **E**, Binding curves of ΔNP-iDr and double phospho mimic mutant ΔNP-IDR S98D T119D titrated to ^15^N labeled KIX measured from ^1^H-^15^N HSQC spectra of the MLL binding pocket. The x-axis represents increasing concentration of IDR or IDR S98D T119D, combined chemical shift differences of KIX residue in the y-axis. **F**, Correlation plot of the combined chemical shift changes of KIX when titrated to IDR 103-107, mutation of the PIM. and ΔNP-IDR at 15 molar equivalents. These results suggest that mutations of the PIM and deletion of NP have similar consequences for the binding of the KIX domain.

To infer if phosphorylation at these sites mediates transactivation, we “simulated” phosphorylation at S98 and T119 by substitutions with aspartic acid (Pearlman et al., 2011; Thorsness and Koshland, 1987). We tested whether S98D and T119D single and double mutants could rescue loss of function due to kinase inhibition described above. We found that substitutions of either S98D or T119D resulted in a significant recovery of transactivation in the DNA-pk kinase inhibition assay. Only the double substitution, however, could completely recover transactivation activity (Figure 6C). In contrast neither phospho-mimicking substitutions rescued inhibition of the kinase GSK-3 (Figure 6D). These results suggest that the phosphorylation of amino acids S98 and T119 are mediated by the kinase DNA-pk.

To test whether the phosphorylation at S98 and T119 affects HOXA11-KIX interaction we performed NMR titration using S98D and T119D substitutions in the ΔNP-IDR construct (ΔNP-IDR S98D/T119D). The residues of KIX that were perturbed upon binding correlated closely with the ΔNP-IDR interaction profile. Interestingly, the binding affinity of the phospho-mimic mutant was almost two times higher (0.19±0.01 mM) than the original ΔNP-IDR (Figure 6E) at the MLL binding site. This result suggests phosphorylation at S98 and T119 sites increases HOXA11’s affinity to CBP.

### The NP interferes with recruitment of CBP to HoxA11

We hypothesized that the NP is interfering with the binding of CBP to the KBD142-146. In order to test this model we compared the combined chemical shift NMR spectra between 1) the KIX domain with ΔNP-IDR (80-152, without the NP), and 2) the KIX domain with IDR (64-152, which contains the NP). Titration of the IDR resulted in very little correlation to the chemical shift changes observed when titrating ΔNP-IDR (80-152) (r^2^=0.25) (Figure S5A). Titration of IDR caused significant perturbation of residues at the N terminus of KIX that were not observed with the ΔNP-IDR construct. These results imply that binding of IDR to KIX is not equivalent to that of ΔNP-IDR, and that the NP alters the interaction of HOXA11 with CBP.

Our mammalian two hybrid results presented above show that the NP could interact with PIM103-107. To understand how the PIM103-107 motif affects KIX binding we mutated it from PGDVL to its ancestral sequence–GDML (IDR 103-107). The combined chemical shift change between the interactions of IDR 103-107 with KIX correlates poorly with that of wild type IDR and KIX (r^2^=0.5) (Figure S5B) and much better to that of ΔNP-IDR and KIX (r^2^=0.82) (Figure 6F). These results suggest that the NP may be interfering with KIX binding by interacting with the hydrophobic residues at PIM 103-107 consistent with our M2H results. The Rosetta models (Figure 5B, C & Figure S5C) identified W73 residue within the NP region as being in close proximity with the α-helical regions. We speculated that W73 might be necessary for mediating the inter-molecular interaction. Indeed, mutation of tryptophan 73 to glycine (W73G) in IDR (IDR W73G) resulted in KIX binding similar to ΔNP-IDR (r^2^=0.82) and IDR103-107 (r^2^=0.90) (Figure S6A&B) supporting our model.

Interestingly titration of ^15^N labeled IDR to unlabeled KIX produced identical chemical shift changes for the same peaks observed during ^15^N labeled ΔNP-IDR titration (Figure S6C). Never the less, we found that these two constructs have distinct binding affinity for KIX with ΔNP-IDR (Kd ≈ 0.04±0.005 mM) exhibiting five times stronger binding than IDR (Kd ≈ 0.2±0.01mM) (Suppl. 6D) consistent with the model that the NP-PIM103 interaction interferes with the recruitment of CBP to HoxA11.

### Changes sufficient for the derived HOXA11-FOXO1 cooperativity

The previous experiments suggest that two evolutionary changes are responsible for the cooperative interaction between HOXA11 and FOXO1: 1) mutations that lead to the derived PIM at AA’s 103-107 and 2) the derived proline residues associated with residues S98 and T119. We introduced these mutations into the AncTh HOXA11 protein and tested their effects in luciferase reporter assays to determine if these mutations are causal for the derived HOXA11-FOXO1 cooperativity. Introducing the derived PIM103 and the proline residues at P97 and P120 were sufficient for giving the ancestral HOXA11 protein activation function (Figure 7A). Interestingly the forward mutated AncTh HOXA11 protein is a FOXO1 independent activator. Only after introducing phosphorylation mimicking residues at S98 and T119 we observed FOXO1 dependent activation (Figure 7B). We conclude that the evolution of the derived protein interaction motif PIM103-107 together with DNA-pk dependent phosphorylation at S98 and T119 is sufficient to cause FOXO1 dependent target gene activation by HOXA11 at the decidual PRL promoter. We note that we do not have a full understanding which amino acid substitutions are necessary to cause the DNA-pk dependent phosphorylation at S98 and T119.

**Figure 7.**
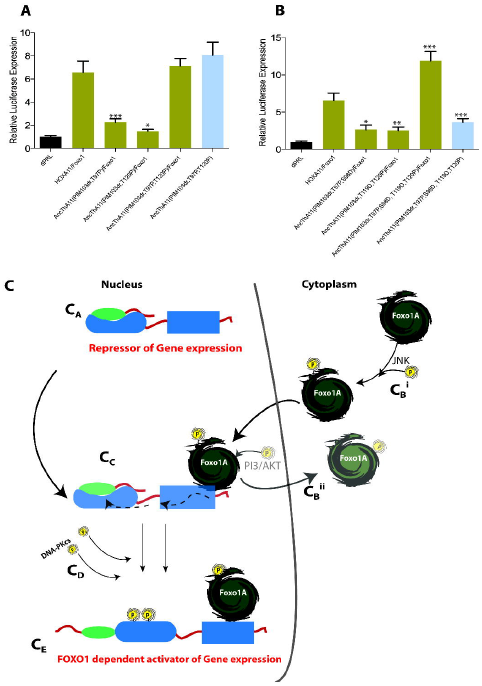
Evolutionary and physiological changes that convert the HOXA11 protein into a transcriptional activator. **A**, Conversion of ancestral therian HOXA11 to a transactivator that is FOXO1 independent. **B**, Conversion of ancestral therian HOXA11 to a FOXO1 dependent activator of luciferase gene expression by mimicking phosphorylation at S98 and T119. Luciferase values are shown as fold changes (mean ± s.e.m., n = 6) relative to the reporter control (dPRL) *p < 0.05, **p < 0.01, ***p < 0.001, ns p > 0.05. **C**, Schematic model illustrating the cooperative regulation of dPRL in decidualized human endometrial stromal cells. **C_A_**, Native HOXA11 is localized in the nucleus and acts as a native transcriptional repressor. **C_B_**, During decidualization, phosphorylation events to the FOXO1 protein is one mechanism, among others, that allow a net increase in nuclear FOXO1. **C_B_^i^**, translocation into the nucleus is facilitated by JNK kinase. **C_B_^ii^**, The retention of nuclear FOXO1 is facilitated by the inhibition of PKB/AKT kinase activity. **C_C_**, Nuclear FOXO1 interaction with HOXA11 possibly induces a structural change (broken black line) that relieves intra-molecular interactions exposing activation. **C_D_**, The recruitment of DNA-pk kinase to S98 and T119 is a derived mechanism of regulating the unmasked activation of HOXA11, by making it a FOXO1 dependent gene regulator. **C_E_**, Foxo1 dependent activated HOXA11 capable of regulating decidual specific genes such as PRL.

## Discussion

At least two TFs, HOXA11 and CEBPB, involved in the regulation of decidual genes have experienced evolutionary changes in their transcriptional activities coincidental with the origin of decidual cells (Lynch et al., 2011; Lynch et al., 2008). In this paper we investigate the derived activity of HOXA11 in response to FOXO1 binding, which evolved in the stem lineage of placental (eutherian) mammals. Using a wide array of experimental approaches, we provide evidence that the derived functional cooperativity between HOXA11 and FOXO1 in placental mammals is due to allosteric regulation of HOXA11 activity by FOXO1 and phosphorylation (summarized in Figure 7C). This work provides evidence for a mechanistic model of how a new context-specific regulatory function of a TF has evolved, while maintaining key features of the ancestral activities. Here we outline a mechanistic model for the evolution of cooperative gene regulation. We then discuss evolutionary changes in the derived HOXA11 protein that enabled the formation of a new mode of gene regulation.

### Ancestral features exploited in the HOXA11/FOXO1 cooperativity

Several key factors necessary for the cooperative gene regulation between HOXA11 and FOXO1 were in place prior to the evolution of functional cooperativity. First, the physical interaction between HOXA11 and FOXO1 evolved prior to the most recent common ancestor of monotremes and humans, i.e. in the stem lineage of all mammals (Brayer et al., 2011). In contrast, the functional cooperativity evolved in the stem lineage of placental mammals (Lynch et al., 2008). The physical interaction between HOXA11 and FOXO1 occurs within the well-conserved homeodomain of the HOXA11 protein (Brayer et al., 2011). By maintaining a high level of constraint within the Homeodomain, the HOXA11 protein retained previously established protein-protein and protein-DNA interactions. We interpret our findings to show that the evolution of a new TF functions can arise from the modification of an already present protein-complex.

Our results demonstrate that the HOXA11 protein has a suppressed ancestral activation function. This activity is masked by an intra-molecular interaction mediated by the newly discovered repressive region, here called NP. This is shown by the fact that a deletion of the NP fragment from the ancestral HOXA11 protein causes FOXO1 independent gene activation (Figure 4A). The HOXA11 proteins from both placental and non-placental mammals are intrinsic repressors if tested in the absence of FOXO1. Only HOXA11 from placental mammals is capable of functionally responding to FOXO1 and unmasking its activation function. This suggests that the role of FOXO1 in the cooperative up-regulation of gene expression in the derived state is to relieve the repression of an already existing activation function. It is likely that HOXA11 is able to interact with other TFs in other cells to unmask its activation potential. If this is the case, the evolutionary event we describe here is an expansion of the set of TFs that HOXA11 can interact to cause transcriptional activation.

### Evolutionary changes in the derived HOXA11

Two kinds of evolutionary changes were identified to be sufficient for converting the ancestral HOXA11 into a context-specific transcriptional activator: i) a derived intra-molecular interaction site and ii) two derived proline substitutions and the phosphorylation of their associated phosphorylation sites.

The first site identified to have a significant impact on the cooperative transactivation was the derived Protein Interaction Motif (PIM, amino acids103-107). The introduction of this site alone, however, could not convert the ancestral HOXA11 into a FOXO1 dependent activator at levels seen with the eutherian HOXA11 (Figure 7A). A back mutation of this site in the eutherian HOXA11 resulted in a FOXO1 independent trans-activator of gene expression (Figure 4D). M2H studies suggest that the derived PIM functions as an interaction site for the repressive region, NP, in the eutherian HOXA11 (Figure 4B). This model is also supported by NMR data.

The second important evolutionary change identified were two proline substitutions at positions T97P and T120P. Individually, these sites in combination with the derived PIM site, in the therian HOXA11, had a significant effect on gene regulation (Figure 7A & B). However, only when both proline substitutions were introduced with the derived PIM site were we able to obtain transactivation at similar levels to the eutherian HOXA11 (Figure 7B).

Further support for the allosteric regulation of transcriptional activation is provided by NMR data. Comparison of chemical shift patterns between ΔNP-IDRL and IDRL with the KIX domain of CBP revealed low similarity in KIX interaction, suggesting that the NP is affecting KIX binding. ΔNP-IDRL binds KIX at the KIX binding motif FDQFF at amino acids 142-146. Finally the tryptophan residue (W) at amino acid position 73 in the NP, as well as mutations at the PIM103-107, leads to a chemical shift pattern very similar to that of ΔNP-IDRL. This result is in agreement with the M2H experiments. Overall our model of allosteric regulation of HoxA11 activity is supported by functional genetic, biochemical (M2H) and NMR evidence.

From these experiments we conclude that the evolution of the derived cooperativity involves at least four mutations. The two threonine ➔ proline substitutions (T97P and T120P) are possible by a single nucleotide substitution at each site. The ancestral mammalian HOXA11 has a glycine residue instead of the derived proline at AA’s 103. A glycine ➔ proline substitution needs a minimum of two nucleotide substitutions. We conclude that at least four nucleotide substitutions are necessary to convert an ancestral HOXA11 into a derived TF able to respond to FOXO1 binding with transcriptional activation.

Multi-functional motifs in disordered regions can allosterically regulate transcriptional activation function. A pertinent example of allosteric regulation is the Ubx TF, where multiple disordered regions control DNA interactions (Liu et al., 2008). Two inhibitory regions (I1 and I2) reduce affinity of Ubx by 2-fold, and by 40-fold, respectively, whereas a regulatory region (R) improves binding in a length-dependent manner. I1 directly contacts ionizable residues of the HD binding interface and also competes with R for the same transient interaction. Such competitive interplay between the disordered regions fine-tunes DNA binding affinity. The allosteric regulation is enabled by the disordered state, which is also maintained in the complex (Fuxreiter et al., 2011). Another recent example is the striking cooperativity switch found with the E1A-CBP-pRb complex (Ferreon et al., 2013).

## Conclusions and perspective

The evolution of TF proteins has been considered unlikely (Carroll, 2005; Prud’homme et al., 2007; Wray, 2007). The rationale was that amino acid substitutions affecting fundamental TF functions would lead to many negative pleiotropic effects. However, research emerging since the late 1990’s has shown that TFs are capable of evolving new functional activities (Wagner and Lynch, 2008). Nevertheless, the mechanisms of how TF function can change are largely unknown.

In this study, we provided evidence for a model of TF evolution where a limited number of amino acid substitutions led to an evolutionary change of the *functional output* of a *pre-existing protein complex.* The derived activation function of the HOXA11::FOXO1 complex evolved through the modulation of a pre-existing intra-molecular repression of a transcriptional activation domain in the HOXA11 protein. The derived mechanism of transcriptional activation has the potential to be context sensitive; it may only happen in cells that express HOXA11 as well as FOXO1 and the appropriate protein kinases. Whether this mode of TF evolution leads to changes in gene regulation limited to certain cell types is an important question to pursue.

## Supplemental Data

Supplemental Data includes Supplemental Data, 6 figures and 4 tables, Material and Methods with any associated references, and can be found with this article online.

## ACKNOWLEDGEMENTS

Financial support has been provided by a grant from the John Templeton Fund (Grant #12793).

